# In vitro testing of flash-frozen sublingual membranes for storage and reproducible permeability studies of macromolecular drugs from solution or nanofiber mats

**DOI:** 10.1101/690990

**Authors:** Pavel Berka, Denisa Stránská, Vladimír Semecký, Karel Berka, Pavel Doležal

## Abstract

Sublingual drug delivery allows systemic delivery of drug without difficulties connected with the gastrointestinal pathway. We developed a new simple protocol for easy-to-use processing and storage of porcine sublingual mucosal membrane for *in vitro* studies using “flash freezing” in liquid nitrogen. All the dextrans used as mucosal membrane integrity and permeability markers permeated only slowly through sublingual mucosa illustrating usability both the “fresh” and “flash frozen” sublingual membranes whereas conventional cold storage “frozen” membranes have shown significantly higher permeabilities for macromolecules due to the sustained damage. The permeability values were too low to expect dextrans to be potential carriers at this context. To test albumin as a drug carrier we compared FITC-albumin permeation from solutions vs. nanofiber mats donors. To increase the amounts and prolong the transport, we manufactured nanofiber mats loaded with fluorescently marked albumin using well-scalable electrospinning technology. Nanofiber mats have allowed albumin passage through the sublingual membrane in similar amounts as from the pure artificial saliva solution. Since salivary washout strictly limits the duration of liquid dosages, nanofiber mats may thus permit prolonged sublingual administration.

**Graphical abstract:** 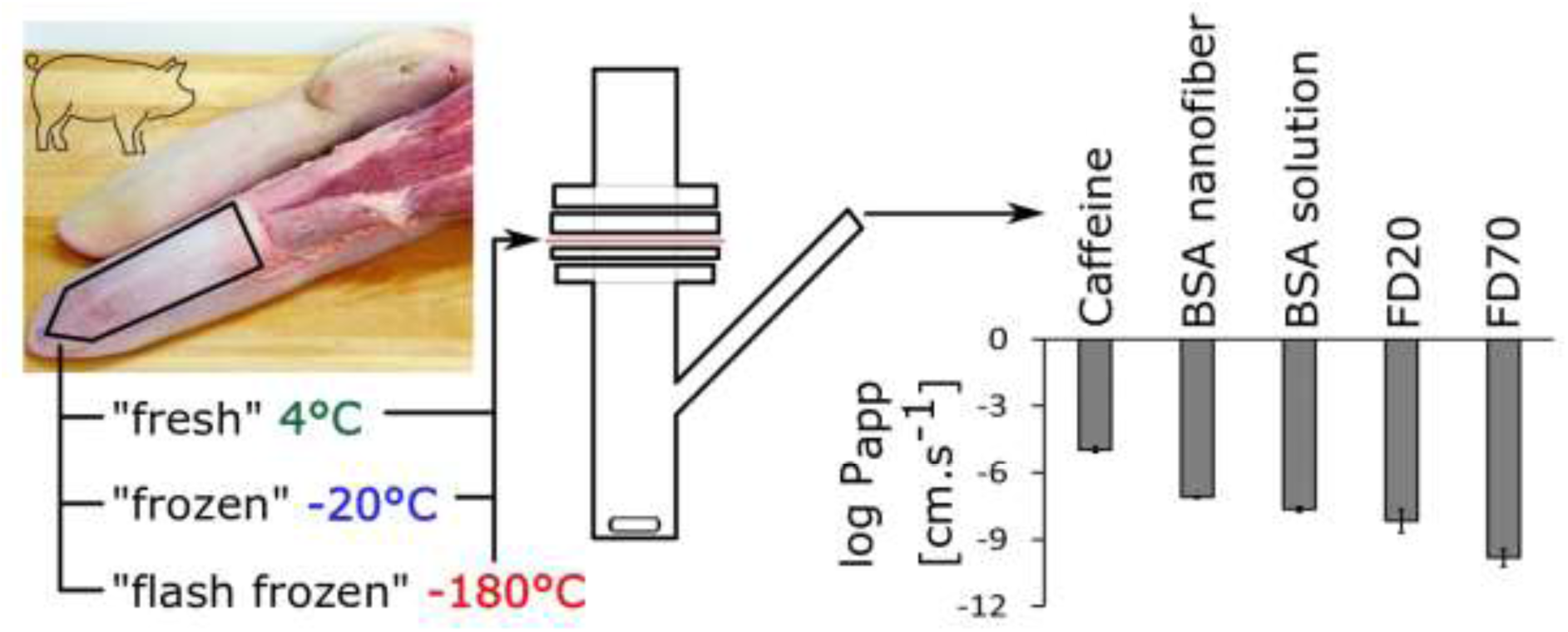

## 1. Introduction

Sublingual and buccal drug systemic administration has been accepted as a potentially efficient alternative route for fast drug delivery or controlled delivery of small molecule drugs for extended periods (Harris and Robinson, 1992) and has been mentioned as a promising route for some biomacromolecules (e.g. peptides, proteins, nucleotides, enzymes, hormones, vaccines) (Castro et al., 2015; Junginger et al., 1999; Kraan et al., 2014). This route has in principle several advantages over conventional oral drug delivery as it avoids the acidic environment and the enzymatic cleavage in the gastrointestinal tract together with hepatic first-pass metabolism. Other typical parenteral routes of administration (i.v, s.c., i.m.) have disadvantages of inherent invasiveness combined with technological problems (mainly due to sterility requirements), poor patient compliance (due to painfulness) as well as price.

The advantages of sublingual and (trans)buccal route include a well accessible route to patients or their carers (Rathbone et al., 2015) for placing or removing several types of medicated preparations (Nicolazzo and Finnin, 2008). Thin and easily permeable non-keratinized oral mucosal areas are perfectly vascularized by blood and lymphatic vessels, only small amounts of enzymes (sublingual amylase, lipase) are present here within close to the pH-neutral environment in the oral cavity; hence the environment does not seriously influence the drug molecule (Veuillez et al., 2001). Moreover, the pre-systemic gut wall and the liver first-pass metabolisms are also avoided (Madhav et al., 2009; Shrestha et al., 2015), and the oral mucosa repairs itself quickly and is not easily damaged (Sattar et al., 2014). A short onset of concentration of unchanged drug in peripheral blood and shortened blood way of the drug to CNS has been proven since 150 years ago (Murrell, 1879).

The disadvantages of oromucosal routes comprise a small area for absorption by passive diffusion resulting in generally small individual doses (usually in milligrams) and low daily drug dosage (a few tens milligrams). Saliva washout, i.e. dilution of the drug by saliva, limits the timespan of the drug in the oral cavity, as it washes out the drug from the application site further into the gastrointestinal tract. Saliva washout thus imposes constraints on the drug dosage forms. (Patel et al., 2011). These disadvantages can be partially overcome by enhancement of absorption using mucoadhesive drug formulation (Montenegro-Nicolini and Morales, 2017; Russo et al., 2016) or chemical enhancers (Madhav et al., 2009).

Reasons for developing oral mucosal drug delivery systems are in detail described in monographies above mentioned (Nicolazzo and Finnin, 2008; Nielsen, 2002; Rathbone et al., 2015) and updated in extensive recent reviews (Brandl and Bauer-Brandl, 2019; Hearnden et al., 2012; Madhav et al., 2009; Mrsny, 2012).

Although a variety of sublingual dosage forms have been described from sublingual tablets or sprays through mucoadhesive wafers, to nanofiber strips (Borbás et al., 2015; Vrbata et al., 2013), and printed films (Wimmer-Teubenbacher et al., 2018), the majority of sublingual drugs (over 20) currently available on the pharmaceutical market are declared as oral thin films. They include for instance opioid pain killers (fentanyl), anti-migraine triptans (sumatriptan), cardiovascular drugs (captopril) and several others (e.g. asenapine, buprenorphine, naloxone, ergoloid, ergotamine, glyceryl trinitrate, isosorbide dinitrate, midazolam, nicotine, prochlorperazine, testosterone, zolpidem), only 3 are intended for delivery of macromolecules (insulin and 2 vaccines) so far, as retrieved from the Micromedex® database (http://www.micromedexsolutions.com). The main reasons are probably linked with the disadvantages mentioned above, namely with the dilution of the drug by saliva followed by its swallowing, without efficient oromucosal absorption.

Nanofiber drug carrier could solve these drawbacks not only for small molecule drugs, as positively shown many times before, e.g. (Thakkar and Misra, 2017), but for at least some macromolecules as well. They can keep a drug at sublingual/buccal absorption place at the maximal concentration for a time sufficient for drug absorption and can prevent the fast dilution and swallowing of the drug, especially when covered by a suitable protective layer. So they possess the better potential to control the conditions within the administration microenvironment then other dosage forms (Morales et al., 2017). Nanofibers are considered as advantageous carriers for drugs belonging to BCS classes II and IV with generally low solubility and problematic bioavailability (Babitha et al., 2017; Paaver et al., 2015). Although these problems and the methods of preparation of the nanofiber drug carriers have been academically discussed for decades, the fundamentally successful technological solution bringing the therapeutic results for these drugs is not at disposal so far. The main reason is that the results are nearly always based on laboratory scale electrospinning techniques and not on well-scalable technology, for instance, such as NanoSpider™ technology (Blakney et al., 2013; Stránská et al., 2019).

The sublingual or buccal permeability of drug is usually tested *in vitro* on porcine mucosal tissue in Franz cells (originally developed for skin permeation studies) as described previously (Nicolazzo and Finnin, 2008; Patel et al., 2012; Sattar et al., 2014). It has similar histological characteristics to the human oral mucosal tissue, and permeability properties are also comparable (Kulkarni et al., 2010), and no other important type of drug transport across sublingual membrane than passive diffusion has been confirmed (Patel et al., 2011). For this reason, it is starting to be considered as a standard in this context. However, fresh mucosa is not always readily available; various procedures for harvesting, treatment and storage have been published and discussed. Comparisons between fresh porcine tissue specimens and those cryoprotected with glycerol 85% for 2 hours and slowly frozen (−1°C/min) to −80 °C and stored in a bio freezer at −80°C revealed no significant effect on permeability for small molecules (Marxen et al., 2016; Sattar et al., 2014).

However, several minutes (about 4 to 6 minutes) of mucosa membrane exposition at 60°C to 65°C during the common cleaning process required by slaughterhouses can damage the sublingual membrane by irreversible changes in proteins and changes (although probably of reversible character) in lipidic structures as well (Mura et al., 2018). These morphology changes might be noticed neither by the eye nor by light microscopy as well as by permeation testing by small molecules, but they could be limiting factor for larger molecules. For this reason, we changed the current slaughterhouse protocol and the tongue region is showered only with lukewarm water (37 °C to 38 °C) to prevent these difficulties. We assumed that this procedure would have less influence not only on proteinaceous epithelial but also all the other associated extracellular structures. We used sodium azide (usual preservation agent against microorganisms) as a fixing agent for sublingual membrane proteins. We also did not try to clear membrane from all the subepithelial tissue, neither mechanically, chemically, enzymatically or thermally. The first mentioned procedure is very time-consuming and may cause mechanical damage, and the other ways are predisposed to change either proteinaceous or lipidic structures of the sublingual membrane as well.

Furthermore, we intentionally did not use any cryoprotectants. They act as intracellular or intercellular cryoprotectants preventing cell damage crystal formation during slow freezing. However, the mechanisms of function of these cryoprotectants show that properties of the sublingual mucosa as a barrier membrane can be changed as well. The influence of freezing on the permeability of buccal mucosa was studied and discussed several times, but the conclusions are not still closed (Marxen et al., 2016). For example, dimethyl sulfoxide (a frequently used cryoprotectant) is linked with pore formation in the cellular membranes (Notman et al., 2006). Other cryoprotectants (e.g. intracellular glycerol, extracellular sucrose, albumin) interact by hydrogen bonding with water and other polar molecules in mucosa as they bond and replace water to prevent ice crystal formation. Therefore, they likely influence intracellular and intercellular spaces within the membrane, which in turn could influence the sublingual permeability.

We need to emphasise that nearly all published data and conclusions are concerned with porcine buccal mucosa. Moreover, the procedures of mucosal membranes treatment described at different experiments were also different. We focused on sublingual nonkeratinized mucosa, and we have known that the problems described for buccal could be very similar but not identical for sublingual mucosa, as the morphological and physiological differences may play a certain role.

The main goal of this study was to synergistically utilise a combination of the properties of sublingual administration route and the nanofibers as a drug carrier. After evaluation of properties of the newly proposed sublingual membrane preparation protocol using solutions of caffeine, and fluorescently labelled dextrans of different molecular weight as neutral polysaccharide molecules. Then, we evaluated *in vitro* permeation of fluorescently labelled albumin released from electrospun-manufactured nanofiber mats using the porcine sublingual membrane and comparing permeability results with a solution of the same polymer in artificial saliva.

## 2. Materials and methods

### 2.1 Materials

Fluorescein isothiocyanate (FITC)-dextrans (FD20 − 16.8 kDa; FD70 − 70 kDa), FITC-albumin, bovine serum albumin (BSA, 66 kDa), eosin Y, formalin and methanol (HPLC grade) were purchased from Sigma Aldrich (Prague, CZ). Caffeine, sodium chloride and xylene were purchased from Dr. Kulich Pharma (Hradec Králové, CZ), haematoxylin from Lachema (Brno, CZ). Polyvinyl alcohol (PVA, type Z 220, the viscosity of 4wt% water solution at 20°C 11.5 – 15 mPas) from Nippon Gohsei (Düsseldorf, GE), polyethylene oxide (PEO, 400 kDa) was purchased from Scientific Polymer Products (New York, USA). Formic acid, phosphoric acid, potassium hydrogen phosphate, potassium dihydrogen phosphate and ethanol (96%) were supplied by Penta Chemicals (Prague, CZ). Sodium azide was purchased from Chemapol (Prague, CZ). Liquid nitrogen was purchased from Linde Gas (Prague, CZ). All the aqueous solutions were prepared with purified water. All the chemicals were used as received without further purification.

Fresh porcine tongues were obtained from the local slaughterhouse Maso Planá (Planá nad Lužnicí, CZ).

### 2.2 Sublingual mucosa membrane preparation

Pieces of mucosa were obtained from the lower side of fresh porcine tongues (*Sus scrofa, var. domestica*) after euthanasia (by the mixture of carbon dioxide with air) by a procedure excluding exposition of mucosa to heat. The intact excised tongues after careful showering with lukewarm water (36°C to 38°C) were put into plastic bags and cooled to 4°C to 6°C in an electric mobile cooling box in which they were transported from a local slaughterhouse to the Faculty laboratory (total time up to 3h).

In the next step, the major muscle mass was cut off. Then, the resulting 2 mm thick mucosal membrane with the remaining muscle was immersed in cold (4°C to 6°C) isotonic phosphate buffer saline (PBS) pH 7.4 containing 0.002% sodium azide for preservation and protein fixation for ca 20 minutes, then further cleaned from most of the muscle and connective tissue by scalpel. The resulting thickness of thus processed membranes was about 0.4 mm to 0.6 mm.

The first part of the obtained membranes (named “fresh” through the manuscript) was then cut into pieces of appropriate size by scissors and used immediately in permeation experiment. The time between porcine euthanasia and the start of the permeation experiment did not exceed 8 hours.

The second part of the membranes (named “flash frozen”) was dried with a cotton tissue, sealed into a plastic bag, then immersed into liquid nitrogen (−180°C) for 60 seconds and stored as “flash frozen” samples in a conventional freezer (−15°C to −25°C) until required.

The third part of the membranes (named “frozen”) was also dried with a cotton tissue and sealed into plastic bags, but without immersing into liquid nitrogen stored in a conventional freezer (−15°C to −25°C) until required.

In a time of permeation experiment, these “fresh”, “flash frozen”, and “frozen” mucosal samples were slowly thawed at 37°C, cut into appropriate pieces (ca. 2 cm × 2 cm) and fixed horizontally between a donor and an acceptor compartment of the Franz diffusion cells. All the mucosa membrane pieces were equilibrated for 15 minutes by acceptor phase after placement into Franz diffusion cells.

### 2.3 *In vitro* permeation

The area exposed for permeation was 1 cm^2^. The phosphate buffer saline (PBS; pH 7.4) was used as an acceptor phase and stirred with a magnetic bar during the experiment. Diffusion cells were placed in a thermostated water bath kept at 37.0 ± 0.5°C.

The permeation testing of dextrans was performed using the donor dispersions in PBS pH 6.8 (0.5 mL) with 0.5% caffeine as a membrane permeability marker with 1% of either FD20 or FD70 dextran. Samples (0.6 mL) of the acceptor phase were withdrawn in pre-determined intervals over 24 hours and replaced with fresh buffer. The samples were briefly stored in a refrigerator until HPLC or spectrofluorimetric determination of the investigated substances was performed with at least three replicates of each measurement. The values presented are calculated as the means with standard deviation (SD) after Dixon Q-test for outlier treatment in MS Excel 2013.

### 2.4 Determination of permeants

The concentration of **caffeine (0.2 kDa)** in the samples of the acceptor phase was assessed using HPLC (Agilent 1200 equipped with UV/VIS and FLD detectors, Agilent, USA). The amount of caffeine was determined using a Zorbax Eclipse plus C18 column (250 x 4.6 mm, 5 μm) and methanol:0.2% aqueous solution of formic acid 1:3 (v/v) as the mobile phase with the flow rate 1.5 mL/min. The UV/VIS detector wavelength was set at 272 nm. The retention time of caffeine peak was 5.75 min.

Spectrofluorimetric quantification of fluorescently labelled **dextrans (FD4, FD20, FD40 and FD70)** in samples of the acceptor phase was carried out on Aminco Bowman Series 2 (ThermoFisher, USA). The excitation wavelength was set at 419 nm and emission wavelength at 529 nm. The system was calibrated using the individual dextran standards dissolved in acceptor buffer pH 7.4.

Spectrofluorometric quantification of fluorescently labelled bovine serine albumin **FITC-BSA (66 kDa)** in samples of the acceptor phase was carried out on Agilent 1200 HPLC with FLD detector using a column bypass (Agilent Technologies, USA). The excitation wavelength was set at 495 nm and emission wavelength at 523 nm. The system was calibrated using standard FITC-BSA solutions in acceptor buffer pH 7.4.

All HPLC data were analysed using Agilent ChemStation software Rev.C.01.06 (Agilent Technologies, USA).

### 2.5 Permeation data treatment

The primary data from spectrofluorimetric or HPLC assay of the samples were further corrected for sampling and replacement of the acceptor phase. The amounts of either FITC-dextrans, FITC-albumin or caffeine which passed through the 1 cm^2^ of sublingual mucosa were obtained. Concentration data were plotted as the cumulative amount of the permeant that diffused from the apical to the serosal side of the epithelium versus time. The apparent permeability coefficient, *P*_app_, was calculated from the formula:

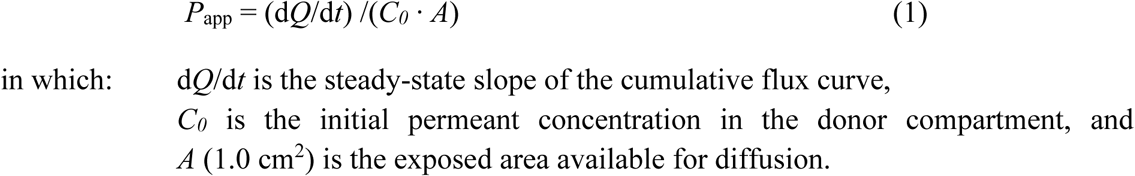

The permeability coefficients were calculated from the linear part and are presented as means ± SD; **n** is given in the pertinent figures. Statistical significance was determined using a t-test, where appropriate.

### 2.6 Integrity testing by impedance measurement

The impedance measurements were performed before and at the end of the permeation tests. The electrical impedance of the sublingual mucosal was recorded using an LCR meter 4080 (Conrad Electronic, Hirschau, Germany) with measuring range 20 Ω–10 MΩ, operated in a parallel mode with an alternating frequency of 120 Hz.

The mucosal samples were mounted into the Franz diffusion cells, the acceptor compartments were filled with PBS at pH 7.4, and the cells were equilibrated at 37°C for 15 min. 500 µL PBS pH 6.8 was poured into the donor compartment. The baseline resistance of the sublingual membrane (Ω⋅cm^−2^) was measured three times in succession at 5-second intervals with two stainless steel wire electrodes inserted into the donor and acceptor compartments of the diffusion cell, respectively.

The buffer solution was removed from the donor compartment using a cotton swab. The solution of the pertinent FITC-dextran as the main expected permeant, also containing 0.5% caffeine as a membrane permeability marker, was then applied to the sublingual membrane.

The second set of impedance data was recorded at the end of the permeation experiment, after 24 h.

### 2.7 Microscopy

Mucosal samples were first thawed at 37°C similarly as in permeation experiments. Then they were placed into 10% formalin, slowly dehydrated in ethanol solutions with increasing concentration (40%, 60%, 85%, 94% ethanol), placed shortly in xylene and embedded in paraffin. Finally, they were cut to 6µm slides by microtome and stained with the haematoxylin-eosin mixture. The obtained slides were examined under an optical microscope Olympus AX70 (Olympus Ltd., Japan) at 100x magnification.

### 2.8 Nanofiber mats preparation

The nanofibrous mats were produced by electrospinning (ES) from polymer solutions using InStrip Technology (www.instar.tech) – a newly adopted method for production of single dose pharmaceutical preparations (Stránská et al., 2019) based on NanoSpider™ technology (Jirsák et al., 2005)

The ES solution consisted of polyvinyl alcohol (PVA), and polyethylene oxide (PEO) in the ratio of 94:6. Both the polymers used were separately dissolved in purified water and then mixed to form a clear, viscous solution. The final concentration of polymers in the solution was 11% (10.3% PVA and 0.7% PEO). Sodium chloride to 0.15% concentration. The mixture of FITC-BSA and BSA (1:1) was added and stirred until a homogenous solution containing 8.5% of the BSA mixture related to the weight of dry matter was obtained and then poured into the container of an ES device. The spinning electrode was in the shape of the wire; electrospinning is nozzle free. The distance between the collector and the spinning electrode was set to 16 cm. After the application of a high voltage (60 kV), nanofibers were formed, drawn up, and then collected on a spunbond textile covering the collector wire electrode. Speed of spunbond movement through the device determined nanofibrous layer thickness (approx. 12.5 g/m^2^).

## 3. Results and Discussion

### 3.1 A new method of membrane preparation

The goal of our study was to assess realistic chances for efficient sublingual permeation of macromolecules either as carriers of small drug molecules or for exerting their own therapeutic or diagnostic effects. First, we had to check the quality of the prepared sublingual membranes. The “fresh” sublingual mucosal membranes obtained from the local slaughterhouse upon further treatment with sodium azide were limited in time to run the tests within 8h post mortem of the animal donors. The same procedure was used for the preparation of “flash frozen” sublingual membranes, but they were additionally rapidly and deeply frozen by liquid nitrogen at −180°C and then kept in ca. −20°C. The same storage was also used for “frozen” sublingual mucosa membranes, but without pre-treatment with liquid nitrogen. No thermal exposition over 38°C and no cryoprotectant were used.

### 3.2 Permeation experiments

The routes of drug molecules across sublingual mucosa have been discussed several times since the 1990s. It is generally accepted that hydrophilic compounds permeate via the paracellular route, while lipophilic drugs would permeate via the transcellular route (Nicolazzo et al., 2003). After sublingual absorption, any hydrophilic molecule must pass a small-distance via extracellular route then across connective tissue and then endothelial layers of blood capillaries or walls of the lymphatic system to reach the systemic circulation. We consider the most bioactive proteins, as well as important drug carriers, including dextrans and albumin, are hydrophilic. Therefore, we use caffeine as a membrane hydrophilic permeability marker.

The results of caffeine permeability show that all the values of *P*_app_ are about of the same order (*P*_app_ ~ 1 100×10^−8^ cm/s), fluctuate in a very similar interval and are not significantly different (p = 0.05) between individual membranes (Fig. 1). Moreover, these results are in good agreement with values found in the literature for sublingual caffeine permeability (*P*_app_ = (1 520 ± 310)×10^−8^ cm/s; Goswami et al., 2013).

**Fig. 1.**
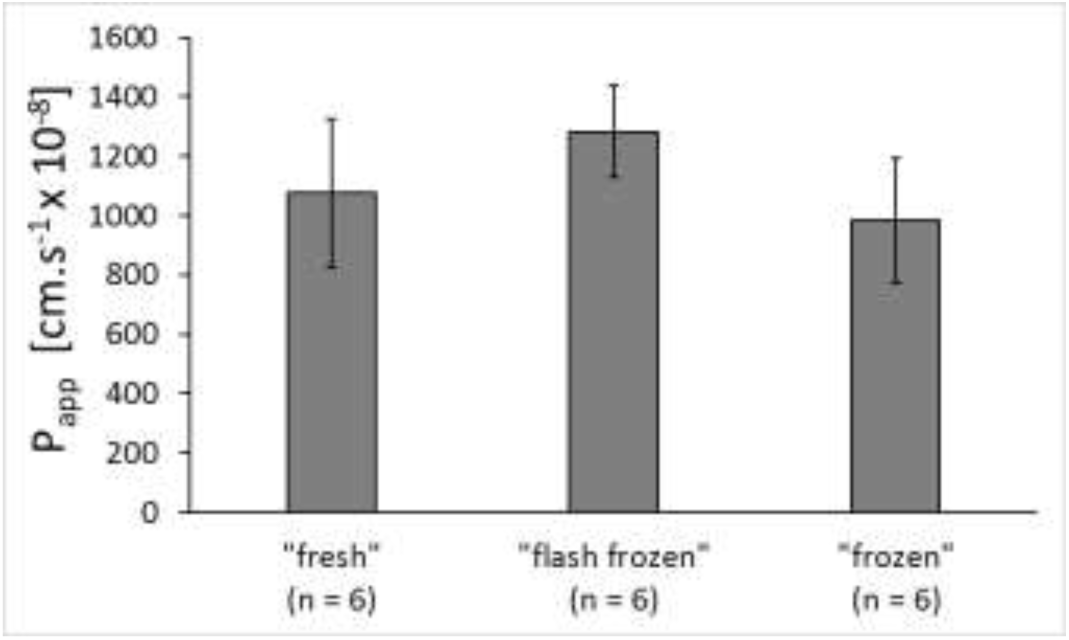
Comparison of caffeine apparent *in vitro* permeability coefficients, *P*_app_, through differently treated porcine sublingual membranes.

Caffeine is a water-soluble substance with no tendency to accumulate within transport routes. As such, it is not able to distinguish among these three differently prepared mucosal membranes. It is worth noting that the similarity of caffeine permeability between fresh and frozen membranes was previously shown also for buccal membranes and in contrast with punctuated membranes (Nicolazzo et al., 2003).

Since the caffeine permeability was remarkably similar in all three membranes, our focus shifted towards the analysis of the permeability of dextrans. First, we have analysed the permeability of FD20 on individual membranes (Fig. 2), and later, we have studied the permeability of other dextrans (Fig. 3).

**Fig. 2.**
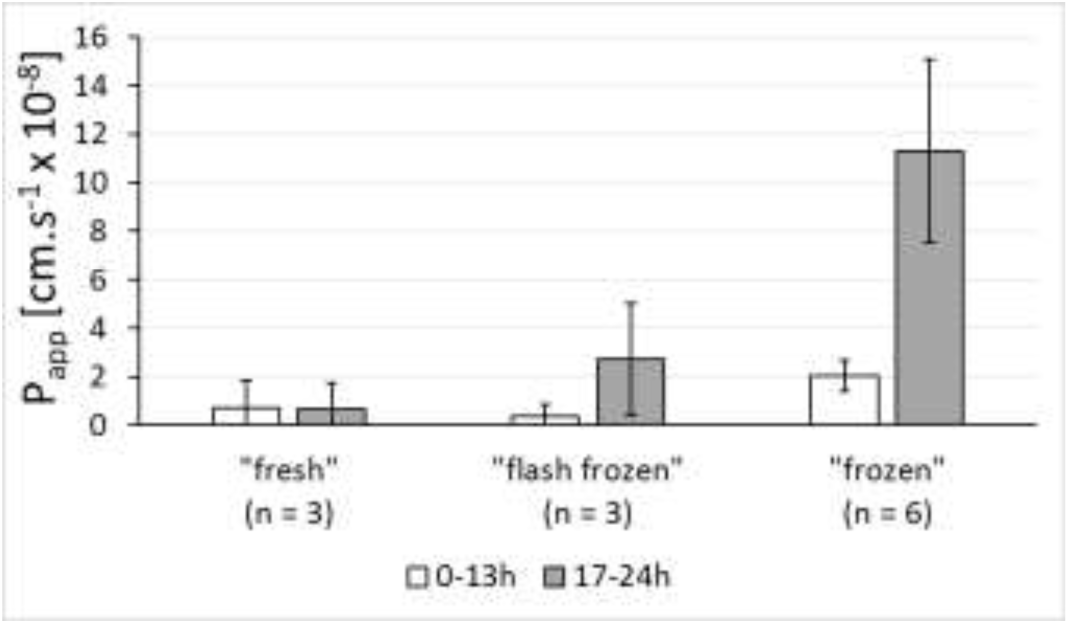
The apparent *in vitro* permeability coefficients, *P*_app_ (± SD), of dextran FD20 through differently treated porcine sublingual membranes up to 13 h and after 17 h of measurement.

**Fig. 3.**
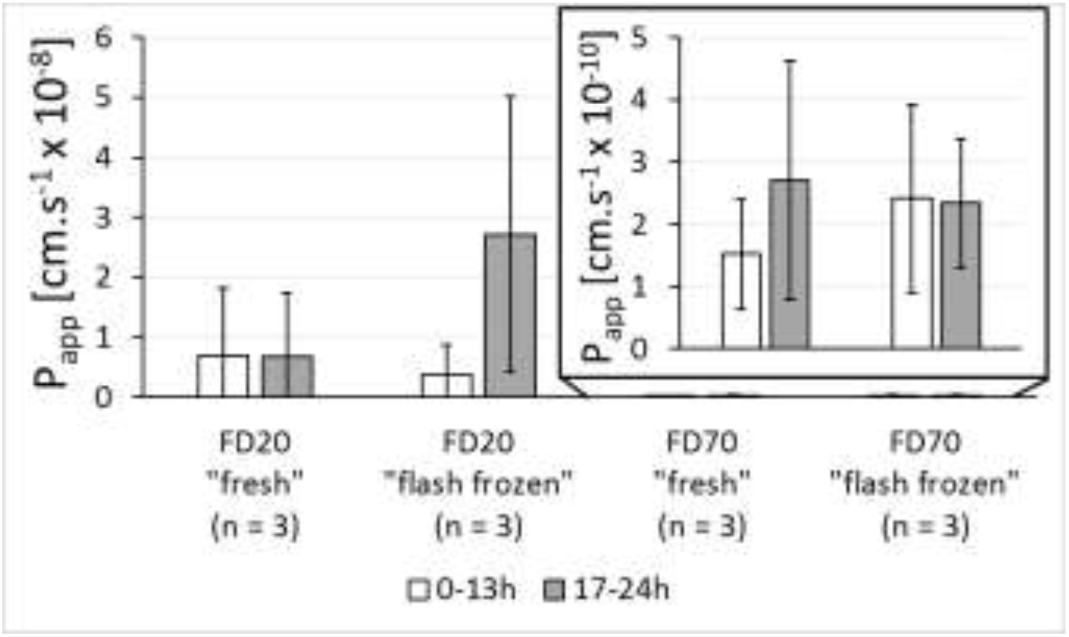
The apparent *in vitro* permeability coefficients, *P*_app_, (± SD) of dextrans FD20 and FD70 through differently treated porcine sublingual membranes up to 13 h and after 17 h of measurement.

FD20 results show high fluctuation of the measured permeability data. Permeability through “fresh” and “flash frozen” mucosal membranes are very similar within the time interval up to 13 h (see Fig. 2, empty columns) from the experiment start. It means that the barrier properties of the “flash frozen” mucosal membranes are probably not influenced neither by the shock freezing by liquid nitrogen nor by the following storage in the conventional freezer at about −20°C. The absence of cryoprotectant seems not to be a problem. The clear differences in barrier properties signalled by increased values of *P*_app_ are visible only after 17 h time interval at “flash frozen” mucosal samples (see Fig. 2, grey column in the middle). Meanwhile, all results obtained for “frozen” mucosal samples show significantly increased permeabilities almost by order of magnitude (without cryoprotectants again).

These results confirm that the “flash frozen” procedure for sublingual membrane processing is a practical easy-to-use preparation method for *in vitro* experiments in sublingual drug delivery studies. It is, however, unlikely that this route of drug administration could be practically associated with a longer time interval than 1 h. At the same time, the FD20 dextran was confirmed as a good integrity marker (Nicolazzo et al., 2003) as its *P*_app_ did not exceed 2⋅10-8 cm.s^−1^ on less damaged sublingual membranes.

Dextrans can be used not only for the membrane permeation or integrity testing but also for the additional aim to estimate molecular weight limiting their passage through the sublingual membrane. The results obtained using the dextrans of higher molecular weight (FD70) are shown on the right-hand side of Fig. 3. Again, the permeability values are similar for both “fresh” and ‘flash frozen” sublingual membranes and they did not significantly differ (p = 0.05) within 13 h interval (see empty columns); however, the values of *P*_app_ are as small as 2⋅10^−10^ cm/s. Interestingly, these values do not significantly increase upon a longer time scale. That means that dextran FD70 is not good integrity marker as it does not differentiate between “fresh” and “flash frozen” sublingual membranes, even as we see that “flash frozen” membrane only showed increased permeation after 17 h probably due to the damage from the freezing/thawing cycle as seen through increased variability between individual samples. Although these data were always obtained only at three replicates, we can conclude that they confirm the usability of the “flash frozen” sublingual membrane for half-day permeability testing of substances up to the molecular sizes like FD70 such as albumin (see Albumin section 3.5 below).

### 3.3 Impedance measurements

To analyse the increased permeability of “frozen” membranes, we have first evaluated its integrity by impedance measurements. At the beginning of the permeation experiment, the mucosa was considered reliable with resistivity higher than 500 Ω⋅cm^−2^. The validity of this resistivity value was evaluated by additional measurement of membranes punctuated with hypodermic needles after the end of the experiment, where the impedance values reached only about 250 Ω⋅cm^−2^.

The impedance values of the “frozen” membranes before the permeability experiment were highly variable among the samples, probably also due to the hand preparation of the samples with a scalpel, where the thickness was not completely uniform. The impedance measurements after the permeability experiment, however, decreased down to the limit indistinguishable from the punctuated membrane (Figure 4). The “frozen” membranes sustained detrimental damage upon the time of the experiment, which explains the increase of the permeability values at the later stages of the experiment.

**Fig. 4.**
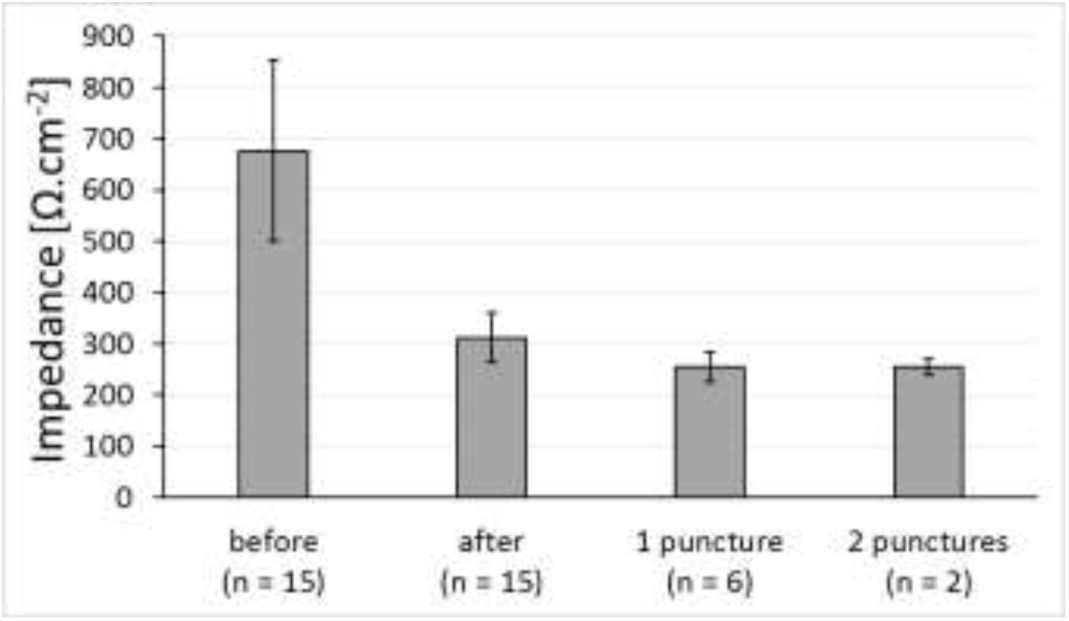
Impedance measurement (±SD) of “frozen” sublingual membranes before and after 24 h of the permeability measurements. The subset of membranes was punctuated afterwards with hypodermic needles to remeasure impedance.

### 3.4 Microscopy

The “frozen” and “flash frozen” sublingual membrane samples were observed by light microscopy just before the permeation experiments (Fig. 5). The left image revealed that the epithelia of "flash frozen" mucosa slices kept intact morphology and integrity. The microphotographs show a stratified, slightly squamous epithelium on the outer (apical) surface with tightly attached cells arranged in regular deeper layers; the papillae are filled with connective material. The surface between epithelial and lamina propria and layer of connective tissue are closely adjacent nearly without bubbles or clefts. Thus, we consider the “flash frozen” membranes as undamaged from the microscopic view.

**Fig. 5.**
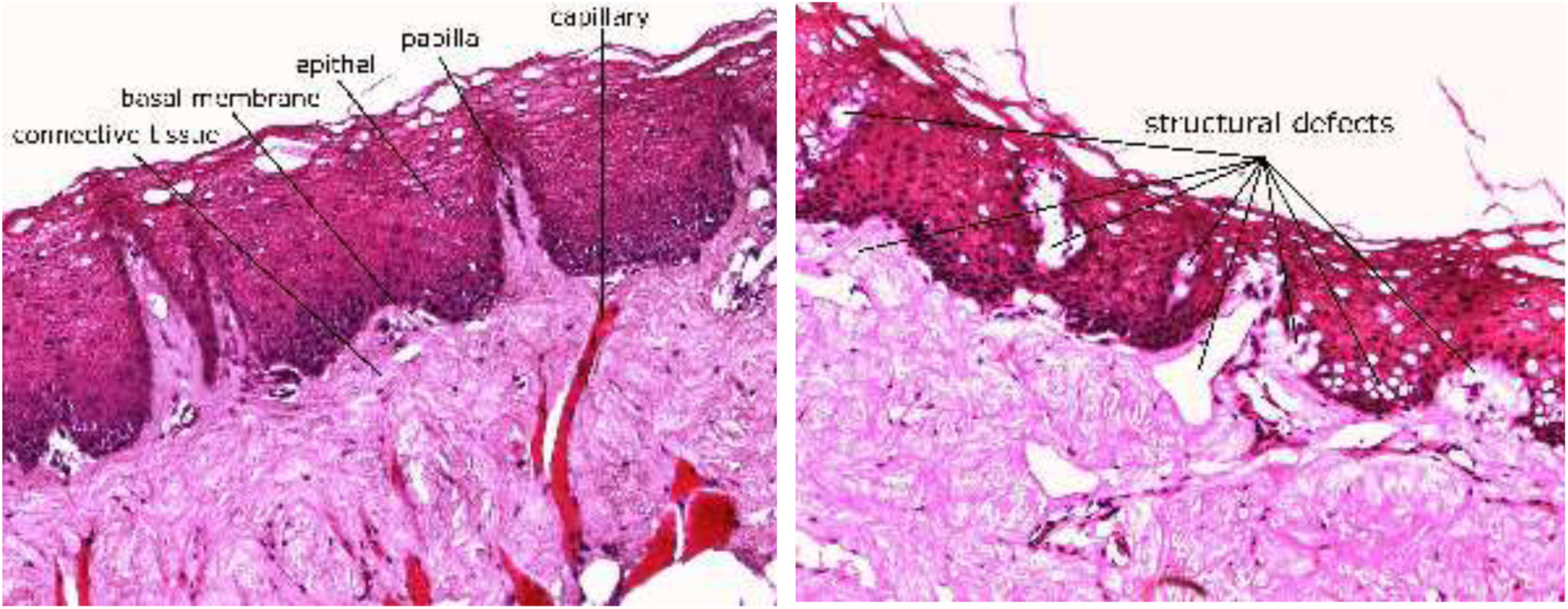
Microscopic slices of thawed “flash frozen” (left) and “frozen” (right) sublingual membranes (100x magnification). “Frozen” sublingual membrane shows multiple structural defects in the lipid content.

On the other hand, “frozen” mucosa slices revealed desquamation and disconnections thorough the sample with void spaces between individual cells. The membrane of such damaged morphology thus cannot serve as a proper barrier, and it explains much higher permeability of the “frozen” sublingual mucosa as revealed by experiments for macromolecular dextrans. Near to the molecular level, the mucosal membrane is probably similarly porous in all samples of different processing history as caffeine permeability was the same in all sublingual membrane preparations.

### 3.5 Albumin

Above mentioned dextrans were used not only for membrane integrity testing but also with an additional aim to estimate molecular weight limiting macromolecular passage through the sublingual membrane. We have tested whether molecular weight is the limiting parameter for the evaluation of permeation of FITC-BSA with similar molecular weight as the FD70 dextran. We have evaluated sublingual permeation of albumin as a protein molecule of molecular weight about 66 kDa with several binding sites for ligands of a different type. It can serve as an almost universal carrier of many small-molecule drugs usable within biological systems. Moreover, it can be considered as a model for the administration of biomacromolecules exerting pharmacological or immunological activity.

However *in vivo* albumin administration through sublingual route in contrast with *in vitro* diffusion cell would require formulation that would not be flown away by salivary washout such as artificial saliva solution. For this reason, we have tried to prepare albumin loaded mats. We have found conditions suitable for the production of albumin loaded nanofibers by using scalable electrospinning technology complying the requirements for pharmaceutically acceptable production for layered nanofiber mats as a part of the sublingual unit dosage forms (see Methods). We have tested whether prepared nanofiber mats can release FITC-BSA and let it pass *in vitro* through the sublingual membrane in a similar fashion as from pure FITC-BSA solution (Fig. 6).

**Fig. 6.**
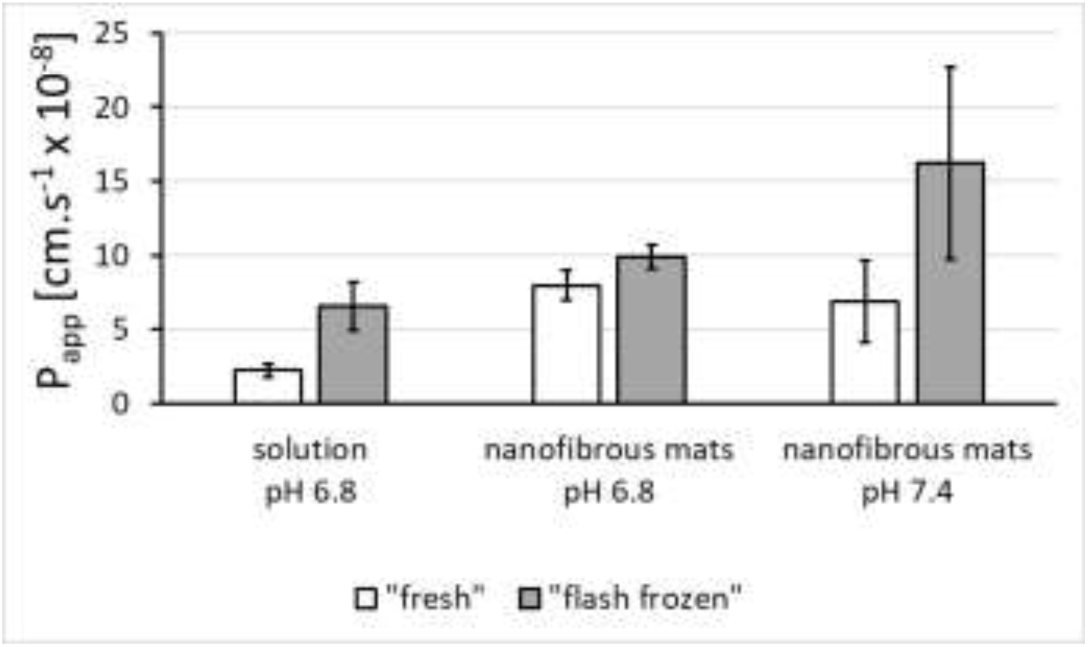
The apparent *in vitro* permeability coefficients, *P*_*app*_ (±SD), of FITC-BSA through differently treated porcine sublingual membranes from solution and from nanofibrous mats during 4h experiment.

FITC-BSA permeated similarly during four-hour long experiment through both “fresh” and “flash frozen” sublingual membranes. Nanofiber-released FITC-BSA permeated approximately two times faster (see Fig. 6 both columns in the middle) than FITC-BSA from solution at pH 6.8, probably due to the tight contact of the nanofibers to the sublingual membrane surface, which increased overall diffusion through the sublingual membrane.

Moreover, it is necessary to stress that used experimental *in vitro* conditions strongly favoured albumin permeation from aqueous solutions in the sense of keeping its concentration at the exposed sublingual membrane area over the whole time of experiment uninfluenced by saliva washout leading to quick decrease of its concentration driving the permeation in seconds and swallowing load drug in minutes. We can emphasise that the situation could be quite different using sublingual mats equipped with a covering nanolayer protecting albumin against saliva washout up for hours.

It is worth notice, that FITC-BSA permeated three orders of magnitude quicker (*P*_*app*_ ~ 1⋅10^−7^ cm/s) than FD70 dextran with comparable molecular weight. The molecular weight is not the only criterion in this context as different types of macromolecules differ in size, shape, compactness, diameter and charge of their molecules, and these parameters also depend on external conditions including the surrounding environment in biological systems. When we compare radius of gyration, *R*_g_, of FITC-BSA at around 32.1 Å (Santos et al., 2003) with FD70 (*R*_g_ ~ 80 Å) and FD20 (*R*_g_ ~ 45 Å), respectively (de Belder, 2003), we can conclude that not molecular weight of polymers but the radius of gyration is clearly a predominant parameter related in the given context.

We can conclude this section, that albumin loaded nanofibers might serve as an interesting carrier option for controlled release of drugs through sublingual route probably even better if used with a suitable cover layer to prevent albumin nanofiber against saliva washout at common physiological conditions in a mouth.

## 4. Conclusions

This study aimed to *in vitro* verify possible sublingual drug administration of model macromolecules using porcine sublingual mucosal membrane. We developed a new simple protocol for easily usable processing and storage of porcine sublingual mucosal membrane for *in vitro* studies using “flash freezing” of the membrane in liquid nitrogen, without any extracellular or intracellular cryoprotectant. Then we estimated the permeability of prepared membranes by in vitro permeation of caffeine on “fresh”, classically “frozen” and “flash frozen” mucosal samples.

While we observed that permeability of caffeine was not affected by mucosal sample preparation in agreement with previous results published by various authors, *in vitro* permeation experiments using FITC-dextrans shown results with significant differences. FITC-dextrans at 20 kDa permeated similarly through “fresh” and “flash frozen” sublingual mucosa at 10^−8^ cm/s rate. In contrast, the conventionally “frozen” membrane was proved to be significantly more permeable for macromolecules and morphologically damaged. Using impedance measurement and light microscopy of stained mucosal samples and by comparing the “flash frozen” and “frozen” mucosa samples, we can conclude that samples conventionally “frozen” at −20°C are are unfit for permeability measurement due to the damage of their epithelial tissue as expected. On the other hand, “flash frozen” membranes pretreated by sodium azide as protein fixative seems to be a useful alternative of “fresh” porcine sublingual mucosa grafts for a half-day *in vitro* permeation testing.

The final *in vitro* permeation experiments using FITC-BSA and “fresh” or “flash frozen” porcine sublingual mucosa confirmed that nanofiber mats allow FITC-BSA to permeate through a sublingual membrane in amounts higher than using its pure solution in artificial saliva even in the steady state conditions without salivary washout.; We can conclude that the amount of FITC-BSA delivered by nanofiber mats might increase the delivered dose of macromolecules.

## Contributions of authors

**Pavel Berka**: Data curation, Formal analysis, Investigation, Methodology, Validation, Visualization, Writing – original draft, Writing – review & editing. **Denisa Stránská**: Investigation, Resources. **Vladimír Semecký**: Investigation, Resources. **Karel Berka**: Conceptualization, Formal analysis, Visualization, Writing – original draft, Writing – review & editing, Funding acquisition. **Pavel Doležal**: Conceptualization, Methodology, Project administration, Resources, Supervision, Writing – original draft, Writing – review & editing, Funding acquisition.

## Conflict of interest

Pavel Berka and Denisa Stránská are employees of InStar Technologies, a.s..

## Supplementary Information

Supplementary Table contains numerical values for all permeabilities reported in the paper.

## Acknowledgements

PB and PD acknowledge support by Charles University Prague project no. *SVV 260 401* and *PRVOUK program 190/11/1110-3.* KB acknowledges support from Czech Science Foundation project no. *17-21122S and ERDF/ESF* Project “Nanotechnologies for Future” project *CZ.02.1.01/0.0/0.0/16_019/0000754*. KB and PB would also like to thank Katka Holá and Štěpánka Berková for immense patience and extreme support.

---

**Supplementary Table 1.**
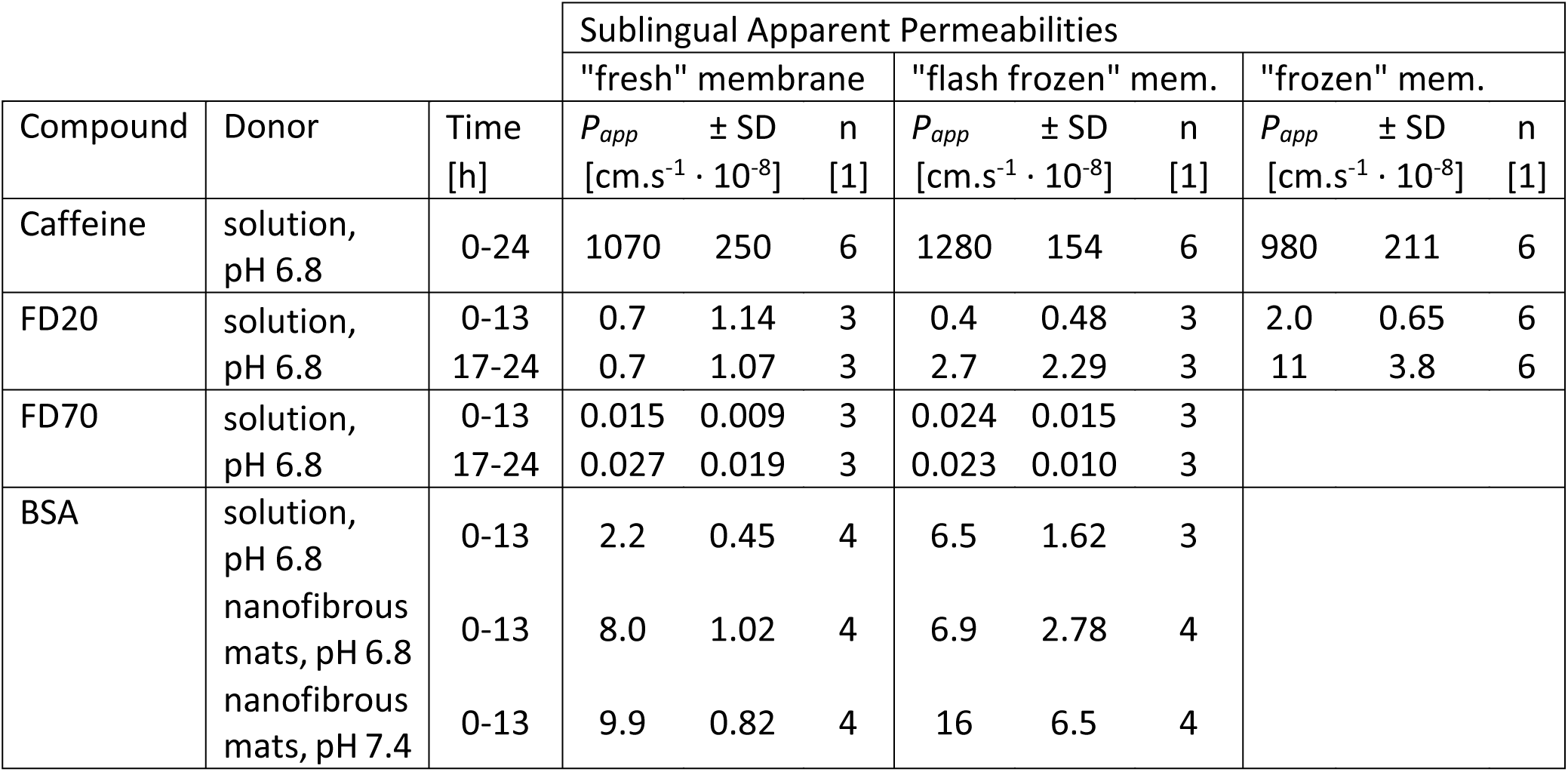
Sublingual Permeabilities Discussed Thorough the Manuscript

## Notes

#### Summary of Updates

We have clarified the manuscript in the descriptions used. We have also added all permeability measurements available in the Supplemental table for easier reuse.

